# High-throughput virus quantification using cytopathic effect area analysis by deep learning

**DOI:** 10.1101/2025.09.11.675495

**Authors:** Michael J. Murphy, Steven Mazur, Elena N. Postnikova, Brett P. Eaton, Gregory A. Kocher, Winston Chu, Matthew G. Lackemeyer, Syed Qasim Gilani, Jens H. Kuhn, Michael R. Holbrook, C. Paul Morris

## Abstract

Traditional infectivity-based virion quantification methods, such as 50% tissue culture infectious dose (TCID_50_) and plaque assays, are typically performed in 6-, 12-, 24-, or 48-well cell culture plates and require manual work and analysis based on legacy protocols. Adaptation of these methods to high-throughput formats (96-, 384- or 1,536-well plates) is challenging due to both assay automation constraints and the lack of well surface area available for reliable analysis. Here, we present a scalable alternative to traditional methods that uses whole-well image thresholding to quantify infectious virions by measuring virus-induced cytopathic effect (CPE) via cell lysis and detachment. The CPE area assay is best positioned as a practical high-throughput preliminary screening tool, effectively quantifying samples within the assay range and flagging samples above or below the concentration thresholds. To improve analysis efficiency and reduce user bias in CPE area selection, we evaluated an nnU-Net model for automated image segmentation against images segmented by manually defined brightness thresholds. The model achieved perfect correlation with manual thresholding (*R^2^*=1.00), showing minimal differences in the identified CPE area and thus validating nnU-Net as a reliable alternative to manual analysis. This approach provides two complementary pipelines: manual thresholding, which tolerates adjustments of hyperparameters, and fully automated segmentation via nnU-Net, which streamlines analysis and enhances throughput. This flexible CPE area assay enables accurate and automated quantification in high-throughput screening formats, thereby greatly accelerating a routine laboratory task while decreasing subjectivity and bias.

**Author summary:** Measuring the amount of virus in a biosample is a critical part of viral research and testing of new treatments. Traditional methods are slow, labor-intensive, and not easily scaled up to handle large numbers of samples. In this study, we developed a new approach that makes the process faster, more efficient, scalable, and less reliant on manual steps. We used microscopy imaging to capture how cells respond to virus infections and analyzed these images automatically to identify and measure areas where the virus had damaged cells. The analysis pipeline allows researchers to fine-tune settings or run the process using machine learning. This method is flexible and adaptable to a variety of viruses and experiments. By increasing the number of samples that can be tested at once while reducing the time and effort needed for analysis, our approach has the potential to accelerate research in virology, drug development, and public health.

## Introduction

Virion quantification methods can be broadly categorized into infectivity assays or component analysis. Infectivity assays strictly measure virions, i.e., virus particles capable of successfully infecting a given host cell, whereas component analysis measures virus proteins, DNA, RNA, or total virus particles [1]. Among infectivity assays, plaque assays have been the gold standard for over 70 years [2–5]. Plaque assays rely on virus-induced cytopathic effect (CPE), which include changes in cell morphology and physiology stemming from virus infection, to produce typically circular clearings of lysed or detached cells, i.e., plaques, within a cell monolayer [2, 3, 6]. Samples are serially diluted to achieve a concentration of virions that produce visibly distinguishable independent plaques, thus facilitating human-annotated counting. A viral titer is then calculated in plaque-forming units (PFU) per volume of sample. Dilutions that generate plaques too numerous to manually count or dilutions with overlapping plaques and unidentifiable epicenters are generally excluded. Requiring a sufficiently large cell monolayer to support the formation of multiple distinct plaques imposes a surface area constraint on the cell culture plate format. Consequently, plaque assays are generally performed in 6-, 12-, 24-, or 48-well plates, which are not conducive to high-throughput, automated platforms.

A widely used alternative for quantifying viral infectivity is an endpoint dilution method known as the 50% tissue culture infectious dose (TCID_50_) assay, which measures CPE in the form of cell lysis and detachment rather than development of discrete plaques [7]. Unlike plaque assays, the TCID_50_ assay is not constrained by surface area limitations, making it more adaptable to high-throughput formats. In practice, the TCID_50_ assay involves exposure of cultured cells to serially diluted pathogen samples, incubation of the cells until CPE becomes visible to the unaided eye, and analysis using statistical methods to calculate the dilution at which 50% of the wells exhibit cell lysis or detachment. Although the TCID_50_ assay can be adapted to 384- and potentially 1,536-well plates, this approach requires many replicates to achieve statistical reliability, which ultimately limits its scalability and throughput.

Traditional plaque and TCID_50_ assays require manual annotation of CPE in cell monolayers to calculate viral titers [2, 6]. Dependence on visual scoring makes both assays labor intensive, prone to bias, and difficult to scale up for high-throughput or automated platforms. Microscopy imaging offers an unbiased solution, replacing manual well inspection with image-based readouts that can be rapidly acquired with many replicates. These large imaging datasets provide a foundation for applying computational approaches. Importantly, microscopy imaging is compatible with high-throughput formats (e.g., 96-, 384-, and 1,536-well plates), enabling experimental output that traditional assays cannot accommodate.

Recent advances in artificial intelligence (AI) have further expanded the potential of microscopy-based assays by enabling automated analysis pipelines [8]. By utilizing convolutional neural networks (CNN) and vision transformers (ViTs), deep learning can reduce or eliminate the need for human annotation through automated plaque counting and CPE quantification [4, 5, 8–11]. However, automated plaque counting remains limited by the surface area needed to distinguish individual plaques, effectively preventing its adaptation to 96-, 384-, and 1,536-well formats. Established automated CPE quantification pipelines based on TCID_50_ continue to rely on binary classification of wells as either CPE-positive or CPE-negative [8]. This approach limits data resolution and requires a large number of replicates to ensure statistically reliable results.

To overcome the limitations of existing assays and methods, we took a stepwise approach that led to an optimized, scalable, and flexible high-throughput alternative to conventional infectivity-based virus quantification assays by utilizing whole-well imaging of cell monolayers to quantify the total area of CPE.

## Results

### Workflow for virus quantification using cytopathic effect area

Permissive cells were seeded into 384-well plates and infected in quadruplicate with either samples of known virus concentration (controls) or experimental samples (Fig 1). Microcrystalline cellulose was used to limit viral diffusion. After a defined incubation period, cell layers were fixed and stained with crystal violet. To image the entire well, the Operetta CLS system acquired nine 10X images per well using both crystal violet fluorescence (excitation: 530–560 nm; emission: 655–760 nm) and brightfield.

To quantify CPE, crystal violet images were segmented by applying a manually defined brightness threshold to distinguish cell monolayer from areas of lysis or detachment. Each pixel was classified as either “signal” (cell monolayer) or “background” (areas of lysis or detachment), depending on whether its intensity falls above or below the chosen threshold. The threshold value was manually selected by visual inspection of the images, often through trial and error, to ensure accurate segmentation. In parallel, brightfield images were processed to define the total well area. The CPE area was then calculated as the total area (from brightfield) minus the area of cell monolayer (from segmented crystal violet).

Controls were arranged as a 24-point dilution gradient across four rows, and standard curves were generated by plotting percent CPE against known viral titers. The equation derived from the standard curve was then applied to experimental samples to estimate viral titers.

**Fig 1.**
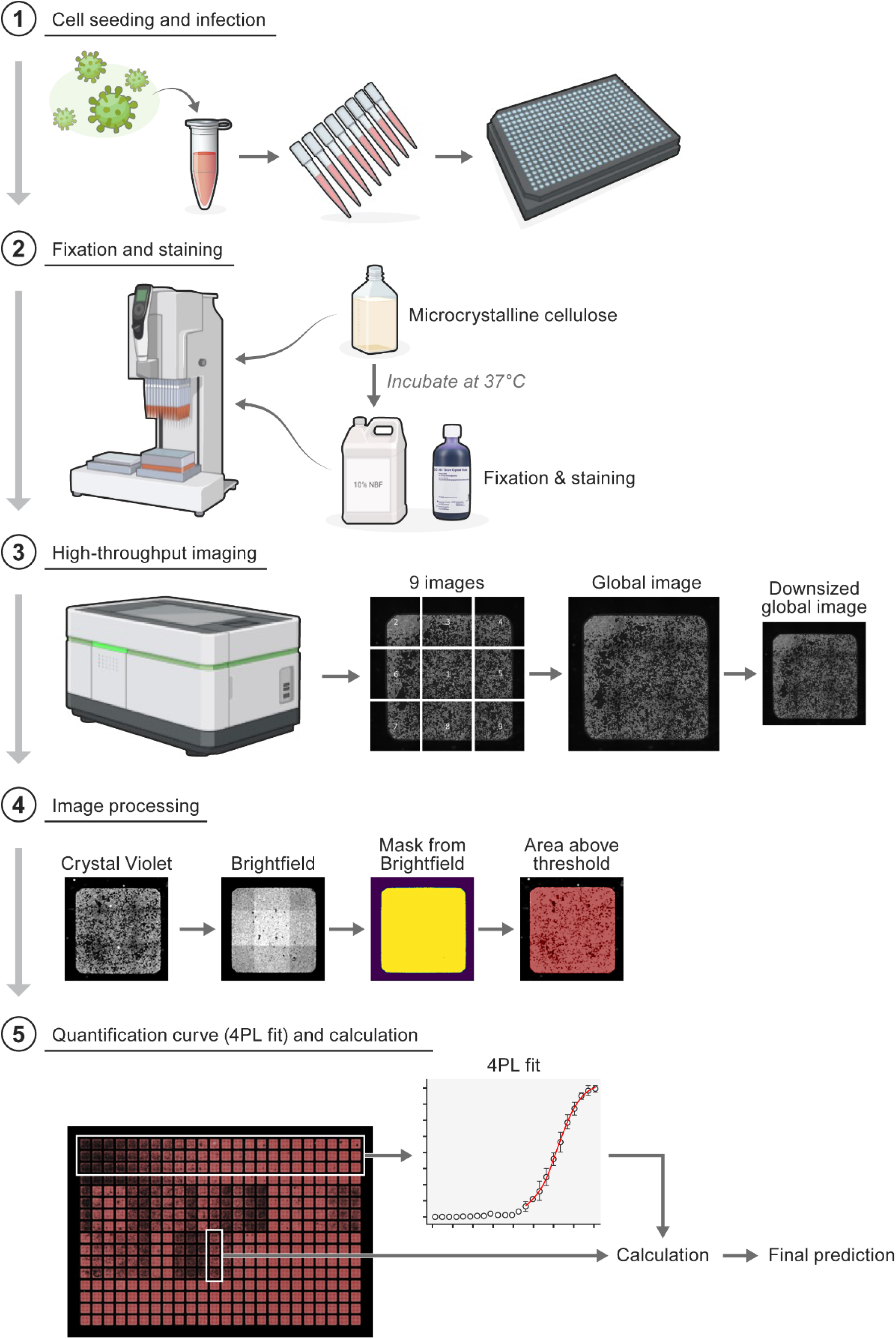
Workflow for virus quantification using cytopathic effect area. **(1)** Suitable virus host cells are grown and exposed to virions. **(2)** Exposed cell monolayers are covered with microcrystalline cellulose, followed by addition of fixation and staining reagents. **(3)** High-throughput microscopy images of stained cell monolayers are acquired and processed. **(4)** Cell monolayers are thresholded using crystal violet and brightfield images. **(5)** Control wells with known viral titers are used to generate standard curves using four-parameter logistic (4PL) regression. The 4PL curve formula is used to predict viral titers in experimental samples.

### Virus infection time influences sensitivity and dynamic range of the cytopathic effect area assay

While optimizing the workflow of the CPE area assay in 384-well plates, we observed that both infection duration and the inherent replication kinetics of the virus strongly influenced how CPE developed over time. To establish proof of principle, we used Rift Valley fever virus (RVFV) vaccine strain under biosafety level 2 (BSL-2) conditions. VeroE6 cells were seeded in 384-well plates and CPE development was assessed at 24, 48, and 72 h post-exposure (**Fig 2**).

At each time point, the percentage of area with discernible CPE increased proportionally with viral titer (**Fig 2A–2C**). This time-dependent relationship illustrates how the assay captures both the extent and progression of infection, confirming its potential for quantitative viral load measurement (**Fig 2D–2F**).

The dynamic range of quantification shifted depending on infection duration. The lower limit of quantification (LLOQ) decreased from an average of 9.6×10^4^ PFU/mL at 24 h to 2.7×10^3^ PFU/mL at 48 h and further to 4.6×10^2^ PFU/mL at 72 h. The upper limit of quantification remained relatively stable, averaging 6.4×10^6^ PFU/mL at 24 and 48 h, with a slight decrease to 4.3×10^6^ PFU/mL at 72 h.

Within the quantifiable range of the assay, predictive performance varied slightly across time points. Predictions were defined as either “hit”, “borderline”, or “miss” based on the fold change (Δ) from the reference titer. A hit was defined as Δ<3.16 (i.e., within a half-log difference), borderline as 3.16<Δ<10 (i.e., between a half-log and full-log difference), and miss as Δ>10 (i.e., greater than a full-log difference). The 24-h time point yielded 86.1% hits, 13.9% borderline, and zero missed predictions in 36 quantified samples (**Fig 2G**). The 48-h time point yielded 79.3% hits, 19.6% borderline, and 1.1% missed predictions in 92 quantified samples (**Fig 2H**). The 72-h time point yielded 85.4% hits, 13.8% borderline, and 0.8% missed predictions in 130 quantified samples (**Fig 2I**). Overall, these findings suggest that infection duration influences the quantifiable range of the assay, with the 48- and 72-h time points proving most suitable for future sample quantification in this study.

**Fig 2.**
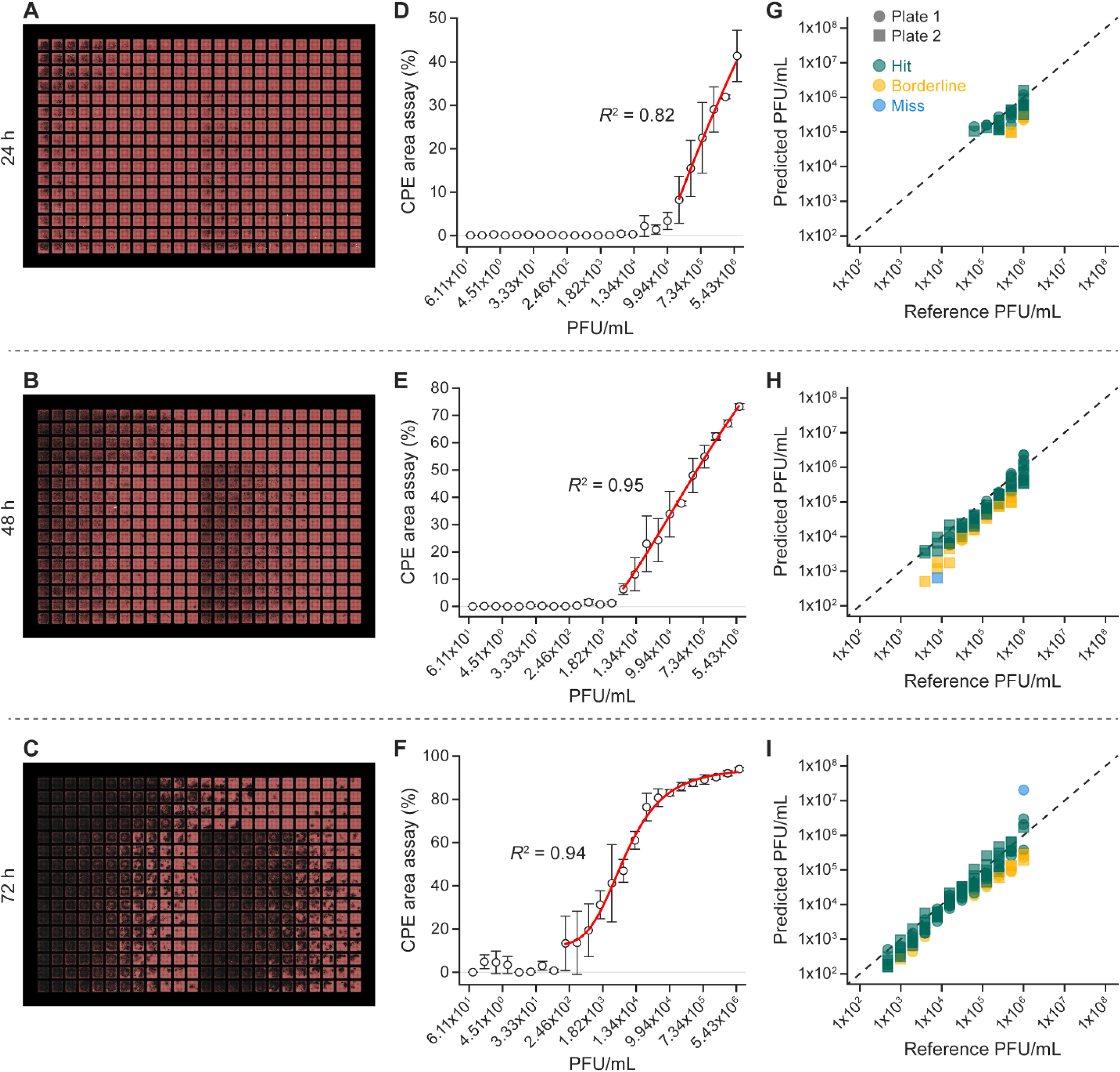
Virus infection time influences sensitivity and dynamic range of the cytopathic effect area assay. **(A, B, C)** Overview images of 384-well Rift Valley fever infection plates at 24, 48, and 72 h. Red coloration indicates crystal-violet-stained cell monolayers. Controls are in rows A–D (all columns, with a decreasing viral titer from left to right columns). Experimental samples are in rows E–P, with replicates split between columns 1–12 and 13–24. **(D, E, F)** Percent CPE areas for controls at each time point with fitted four-parameter logistic (4PL) regression curves. **(G, H, I)** Predictions generated by applying the standard curve formula to percent CPE areas of experimental samples. Plate 1 and Plate 2 represent biological replicates. Predictions were defined as either “hit”, “borderline”, or “miss” based on the fold change (Δ) from the reference titer. A hit was defined as Δ<3.16 (i.e., within a half-log difference), borderline as 3.16<Δ<10 (i.e., between a half-log and full-log difference), and miss as Δ>10 (i.e., greater than a full-log difference).

### High-throughput cytopathic effect area assay provides a scalable alternative to traditional plaque assays

After establishing optimal infection conditions for RVFV in the CPE area assay, we directly compared assay performance at the 48-h time point with the traditional six-well plaque assay using 40 experimental samples (**Fig 3**). The standard curve for the CPE area assay was generated from a 24-point, 2-fold dilution series spanning from 7.63x10^-1^ to 6.40x10^6^ PFU/mL. The 40 experimental samples ranged in concentration from 4.12x10^2^ to 6.4x10^6^ PFU/mL.

All samples were quantifiable using the plaque assay, whereas approximately 30% of the lowest-titer samples were below the LLOQ in the CPE area assay. The plaque assay was highly accurate, with 97.5% of quantified samples falling within a full-log and 95% within a half-log of the reference titers. The 384-well CPE area assay was somewhat variable across plates but maintained good overall accuracy, quantifying 100% of samples within a full-log and 87.5% of samples within a half-log of the reference titers. Within its established limits of quantification, the CPE area assay produced results consistent with the traditional plaque assay, demonstrating its potential as a high-throughput preliminary screening tool.

**Fig 3.**
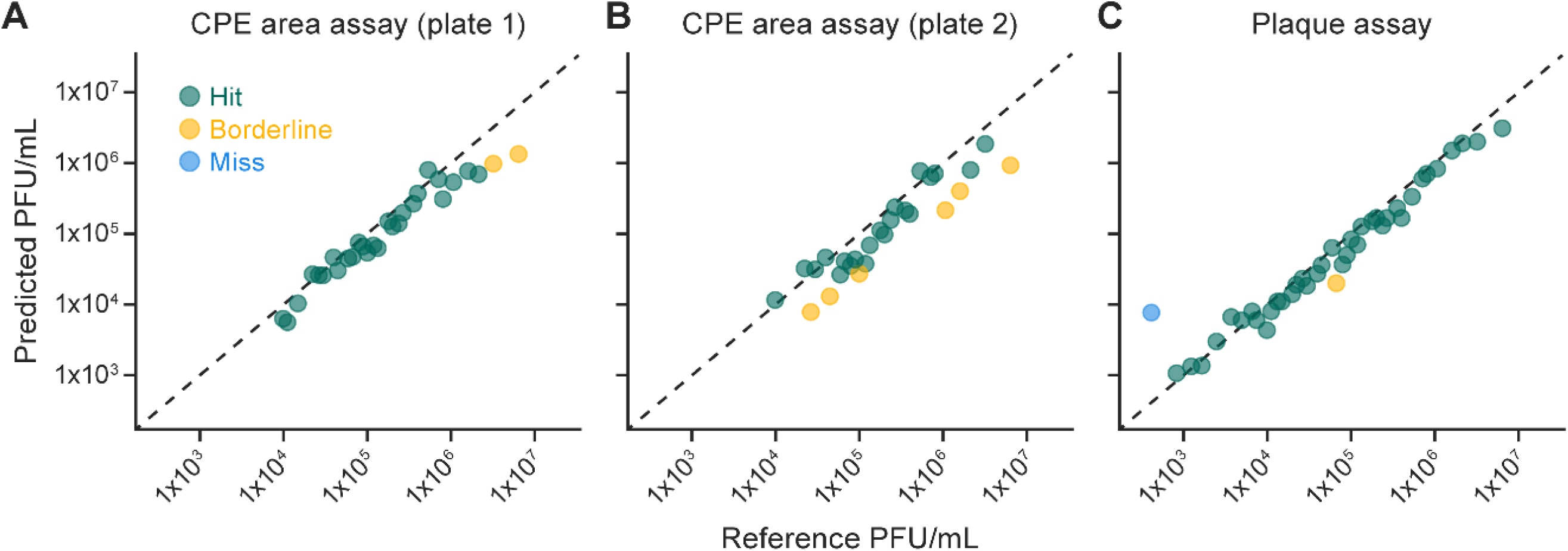
High-throughput cytopathic effect area assay provides a scalable alternative to traditional plaque assays. Comparison of the accuracy and dynamic range of the CPE area assay versus the traditional plaque assay using 40 experimental samples. Predictions were defined as either “hit”, “borderline”, or “miss” based on the fold change (Δ) from the reference titer. A hit was defined as Δ<3.16 (i.e., within a half-log difference), borderline as 3.16<Δ<10 (i.e., between a half-log and full-log difference), and miss as Δ>10 (i.e., greater than a full-log difference). **(A)** CPE area assay predictions compared to reference titers for experimental samples in plate 1. **(B)** CPE area assay predictions compared to reference titers for experimental samples in plate 2 (biological replicate). **(C)** Traditional plaque assay predictions compared to reference titers for experimental samples.

### Cytopathic effect area assay with manual thresholding demonstrates greater accuracy and range than deep learning approaches

During optimization of the CPE area assay with RVFV, we observed distinct spatial patterns of CPE that varied with viral titers. We hypothesized that these spatial features could provide additional information about infection kinetics beyond the area of CPE alone and that deep learning models could incorporate this information to more accurately predict viral titer.

To test this, we trained a Siamese neural network (SNN) and fine-tuned a pretrained ViT model, with the manually thresholded CPE area assay serving as the baseline method for comparison (**Fig 4A**). The SNN was chosen because CNNs are well-established in medical imaging, and the Siamese framework offers the potential to reduce the number of controls required for a standard curve[12, 13]. The SNN consisted of two parallel CNNs based on the AlexNet architecture [14], sharing weights and trained with a contrastive loss function to measure similarity between image feature representations of a sample image and an “anchor” image with a known viral titer (**Fig 4B**).

We also compared the CPE area assay against a ViT model, as recent image classification benchmarks have demonstrated that ViTs can outperform CNNs in image classification tasks, especially with large pretraining datasets [15]. To overcome the limited size of our dataset, we fine-tuned an ImageNet-pretrained ViT-B/16 using the same training data as the SNN. This approach enabled us to leverage the ViT’s ability to capture long-range dependencies and high-level spatial features without requiring extensive infection image data (**Fig 4C**)[15].

For training, 3,840 wells were exposed to RVFV using a broad range of viral titers. Full monolayer images were collected of each of the wells. To evaluate model performance, 384-well plates fixed at 48 and 72 h post-exposure were analyzed, with controls arranged in rows A–D as a 24-point, 2-fold dilution series spanning from 7.63x10^-1^ to 6.4 x 10^6^ PFU/mL. Experimental samples were placed in rows E–P, with replicates distributed across columns 1–12 and 13–24, ranging in concentration from 4.12x10^2^ to 6.4x10^6^ PFU/mL.

**Fig 4.**
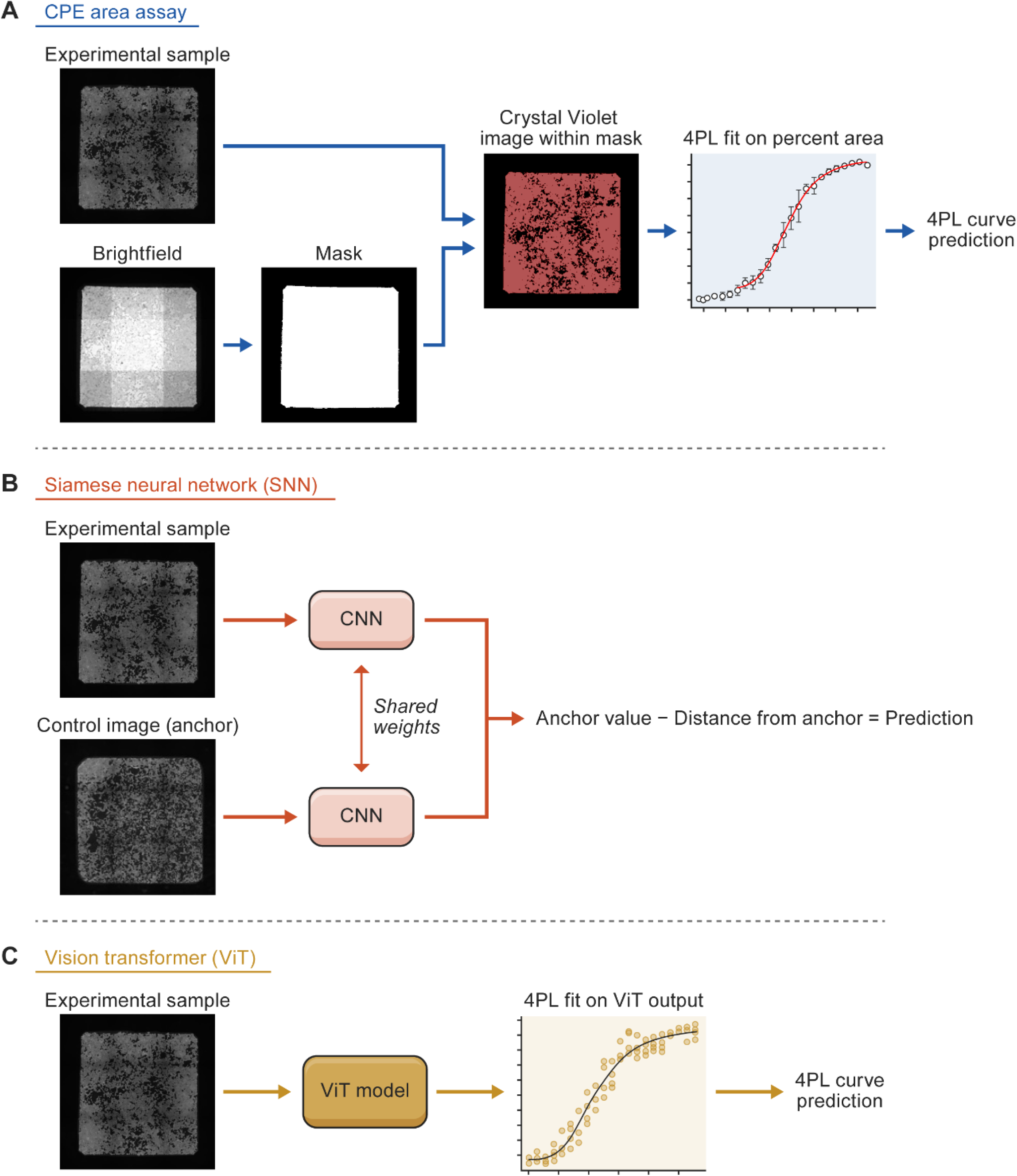
Workflow comparison for cytopathic effect area analysis via thresholding, vision transformer (ViT), and Siamese neural network (SNN). **(A)** CPE area assay method. Brightfield images are used to segment the full area of each well, generating a mask that defines the region of interest. Crystal violet images are then thresholded within this mask. Control wells with known viral titers are used to generate a standard curve via four-parameter logistic (4PL) regression, which is then used to predict viral titers in experimental samples. **(B)** SNN method. A crystal violet image from a control well with a known virus concentration is loaded as the anchor image. An experimental sample image is loaded in parallel, and both images are passed through identical convolutional neural networks (CNNs). The CNNs were trained on images from 3,840 RVFV-exposed independent cell monolayers. The network predicts a distance between the image embeddings, which is subtracted from the known concentration of the anchor to estimate the virus concentration of the experimental sample. **(C)** ViT method. Crystal violet images are passed through an ImageNet-pretrained ViT-B/16 that was fine-tuned on the same RVFV training dataset as the SNN. The output is a single value associated with each image. Values from control wells are used to generate a standard curve via 4PL regression, which is then used to predict virus concentrations in experimental samples.

The dynamic ranges of the CPE area assay and the ViT model were broadly similar (**Fig 5**). At 48 h after virus exposure, the LLOQ was 5.15×10^3^ PFU/mL for the CPE area assay and 7.81×10^2^ PFU/mL for the ViT. By 72 h, the LLOQ extended to 5.45×10^1^ PFU/mL for the CPE area assay and 1.95×10^2^ PFU/mL for the ViT. In contrast, the SNN performed poorly at both time points (**Fig 5C**) despite performing strongly on the original test dataset following training (**Fig S1**).

At 48 h after virus exposure, the CPE area assay identified 36 predictions as hits, five as borderline, and zero as misses (**Fig 5A**). The ViT model identified 32 predictions as hits, 24 as borderline, and seven as misses (**Fig 5B**). When samples accurately identified as below the limit of detection were also classified as hits, the CPE area assay yielded 67 predictions as hits, five as borderline, and zero as misses. This equates to 100% accuracy within 1 log and 93% accuracy within ½ log. Under the same criteria, the ViT model yielded 38 predictions as hits, 26 as borderline, and eight as misses, corresponding to 88% accuracy within 1 log and 53% accuracy within ½ log (**Fig 5D**).

At 72 h after virus exposure, the CPE area assay again performed strongly, yielding 65 predictions as hits, seven as borderline, and zero as misses. The highest accuracy was observed in mid-range and lower concentration samples, with reduced precision at higher concentrations. This decline may be attributed to saturation, suggesting that diluting high-titer samples could improve prediction accuracy in this range. The ViT model produced similar results, with 63 predictions as hits, nine as borderline, and zero as misses. Like the CPE method, the ViT model’s highest accuracy was in the mid-to-low concentration range, with diminishing precision at higher concentrations. Overall, these results show that the CPE area assay outperforms the deep learning models in both accuracy and dynamic range.

**Fig 5.**
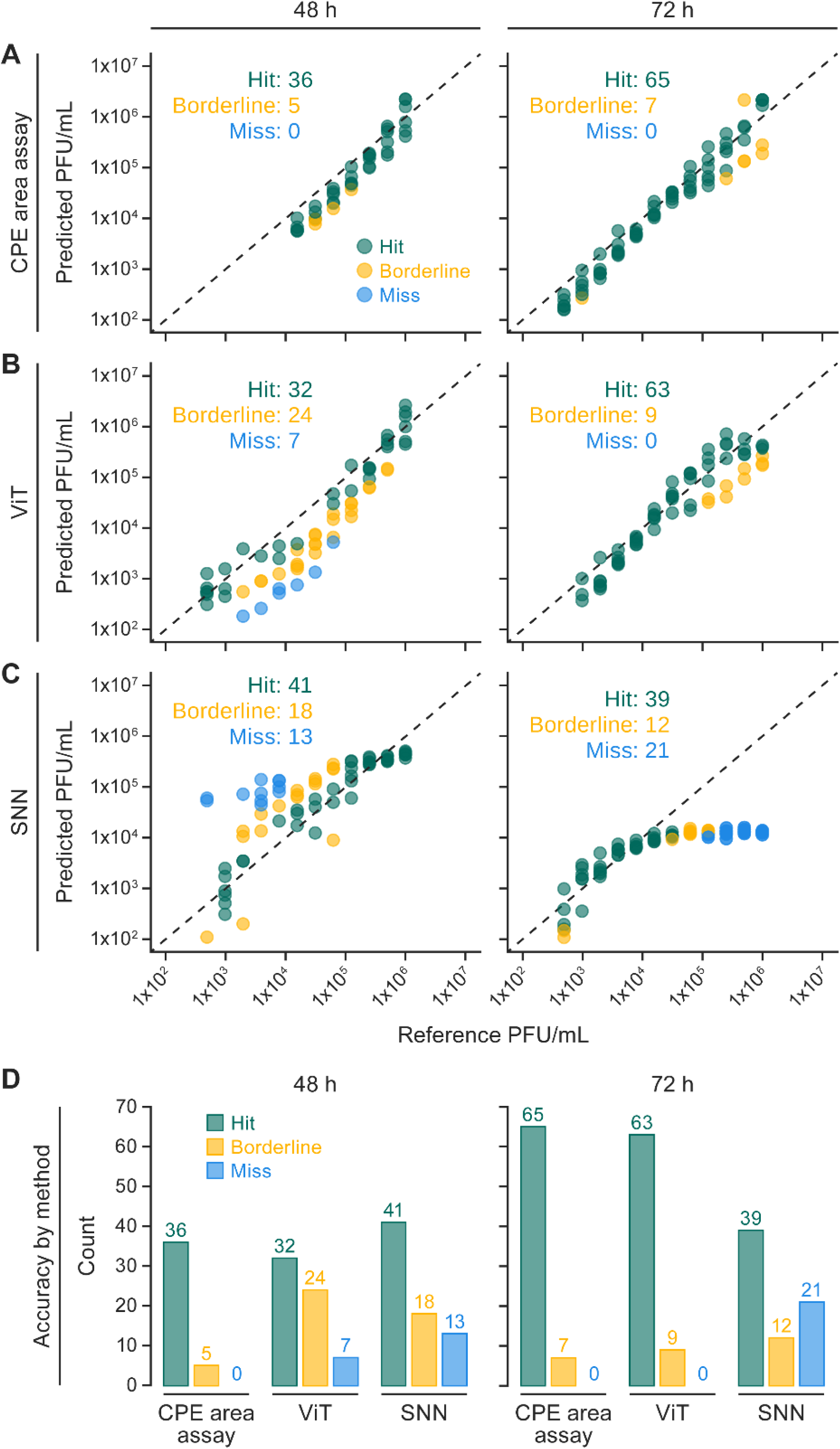
Cytopathic effect area assay predicts viral titer more accurately than the vision transformer or Siamese neural network models. Comparison of the accuracy and dynamic range of the cytopathic effect (CPE) area assay using manual thresholds versus deep learning methods using 72 experimental samples. Predictions were defined as either “hit”, “borderline”, or “miss”, based on the fold change (Δ) from the reference titer. A hit was defined as Δ<3.16 (i.e., within a half-log difference), borderline as 3.16<Δ<10 (i.e., between a half-log and full-log difference), and miss as Δ>10 (i.e., greater than a full-log difference). **(A)** CPE area assay by manual threshold, **(B)** vision transformer (ViT) model, and **(C)** Siamese neural network (SNN) model predictions at 48 and 72 h after virus exposure. **(D)** Accuracy of each method, accounting for predictions both above and below the established limits of quantification.

### nnU-Net automated segmentation is a replacement for manual thresholding in CPE area identification

While most of the CPE area assay with manual thresholding is automated, it does require manual selection of crystal violet and brightfield intensity thresholds, which then determine the areas of plaque and cell monolayer. To improve consistency, remove bias, and decrease labor in this assay, we trained a nnU-Net framework on thresholded images to automatically segment the whole-well area from brightfield images and the cell monolayer from crystal violet images without manual thresholding (**Fig 6**). nnU-Net is a self-configuring deep learning semantic segmentation framework and has demonstrated robust performance in various biomedical image segmentation tasks without the need for manual tuning of hyperparameters [16]. When applied to a test plate not used during training or validation, the model produced segmentation masks that closely matched the thresholded images, with only minor pixel-level differences (**Fig 6A**). Quantification of the CPE area showed excellent concordance between the two methods, with an *R^2^* of 1.00 (**Fig 6B**), indicating that the nnU-Net model can replicate manual thresholding with high accuracy.

**Fig 6.**
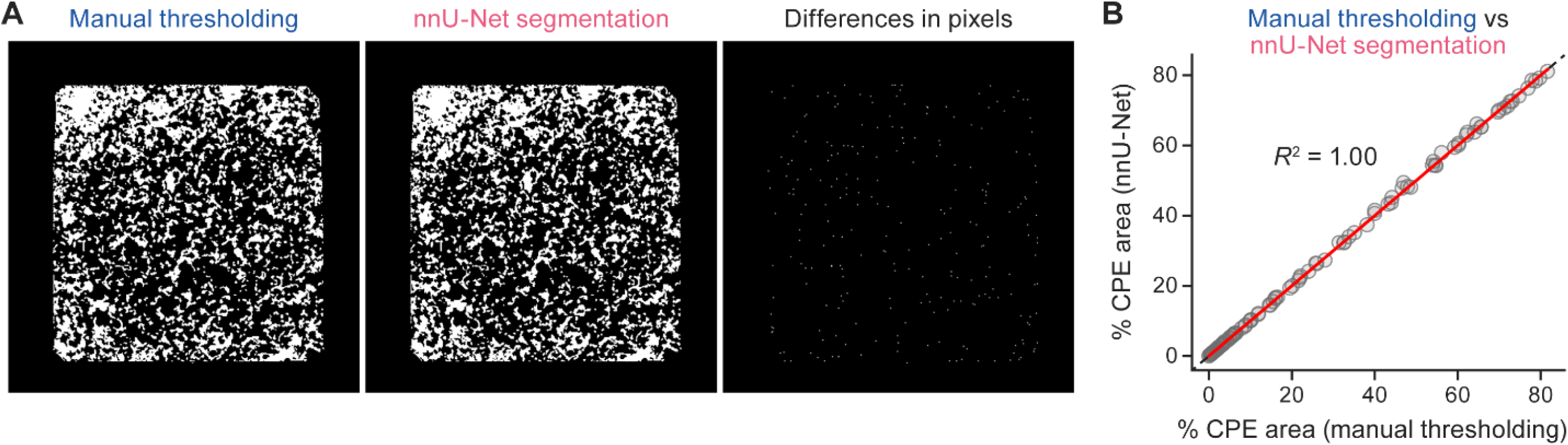
nnU-Net automated segmentation as a replacement for manual thresholding for CPE area identification. Pixel differences between manual thresholding and nnU-Net segmentation identification of monolayer. **(A)** An example image using segmentation from a manually determined threshold value and automated segmentation from nnU-Net. **(B)** Percent cytopathic effect (CPE) area calculated from manually thresholded images compared to percent CPE area calculated by automated segmentation from nnU-Net.

To evaluate the utility of the CPE area assay and demonstrate its application in maximum (biosafety level 4 [BSL-4]) containment laboratory settings, the assay was evaluated for use with eastern equine encephalitis virus (EEEV), Nipah virus (NiV), and severe acute respiratory syndrome coronavirus 2 (SARS-CoV-2). Although EEEV and SARS-CoV-2 are not classified as BSL-4 pathogens, all experiments were performed under uniform maximum-containment procedures to ensure consistent conditions across viruses. For each virus, results from manual image thresholding were compared to automated segmentation outputs generated by the nnU-Net model. This comparison provided further validation of the model’s ability to generalize across distinct CPE morphologies and varying assay conditions (**Fig 7**).

These two methods gave a medium-sized assay range for EEEV (8.34×10^2^–5.48×10^6^ PFU/mL for manual thresholding and 6.97×10^2^–5.43×10^6^ PFU/mL for nnU-Net segmentation), with smaller ranges for NiV (1.38×10^3^–7.72×10^5^ PFU/mL for manual thresholding and 5.25×10^3^–2.72×10^6^ PFU/mL for nnU-Net segmentation) and RVFV (5.63×10^3^–3.39×10^6^ PFU/mL for manual thresholding and 5.25×10^3^–2.72×10^6^ PFU/mL for nnU-Net segmentation). The range for SARS-CoV-2 was exceptional (8.71×10^1^–4.04×10^8^ PFU/mL for manual thresholding and 8.20×10^1^–4.83×10^8^ PFU/mL for nnU-Net segmentation). These methods both yielded very accurate results, predicting greater than 99% of samples within 1 log and approximately 77% within ½ log for all viruses.

**Fig 7.**
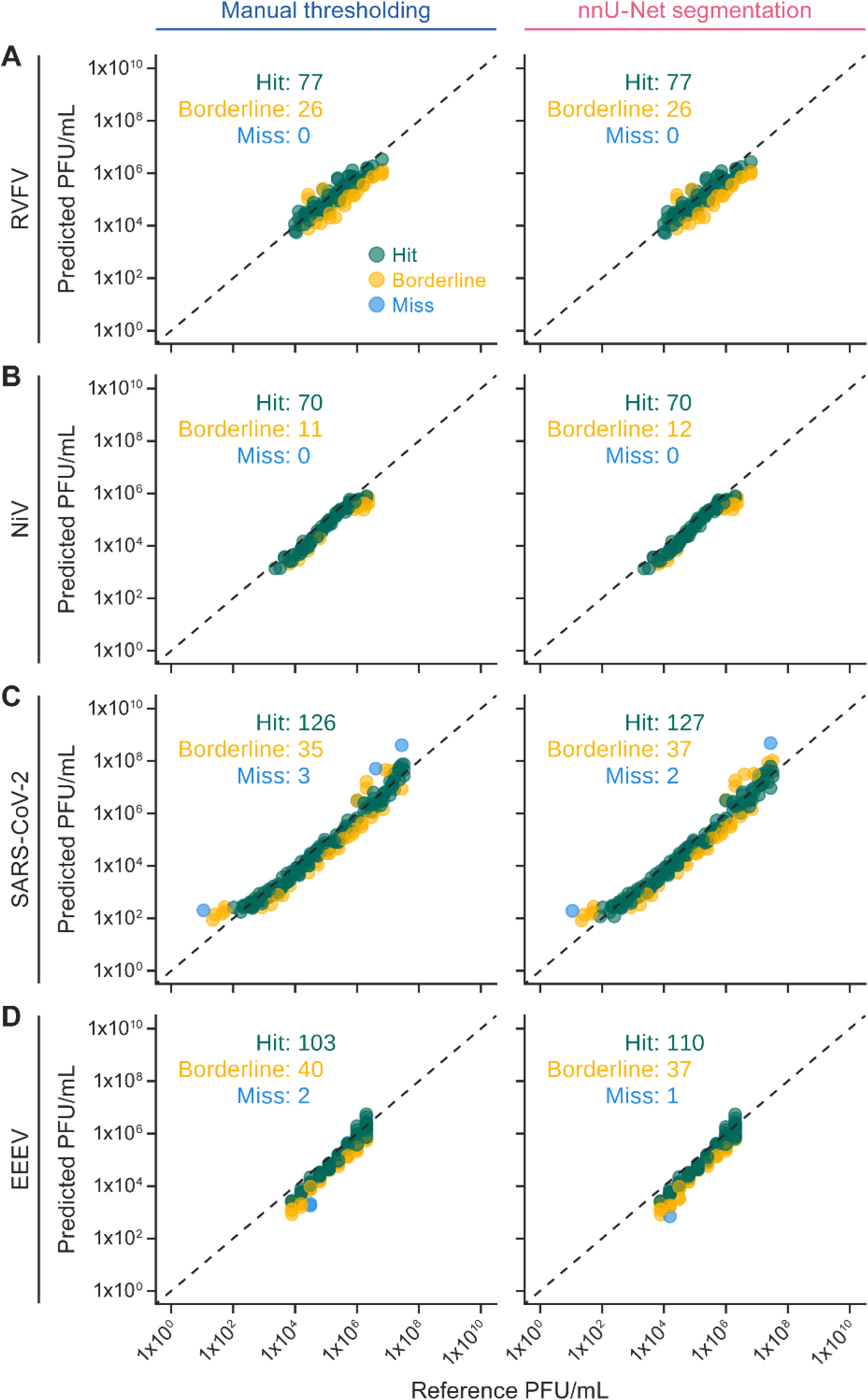
Cytopathic effect area using manual thresholding or nnU-Net segmentation can accurately predict viral titer across multiple viruses. Comparison of the accuracy and dynamic range of cytopathic effect (CPE) area assays using manually thresholded images and nnU-Net automated segmentation. Predictions were defined as either “hit”, “borderline”, or “miss” based on the fold change (Δ) from the reference titer. A hit was defined as Δ<3.16 (i.e., within a half-log difference), borderline as 3.16<Δ<10 (i.e., between a half-log and full-log difference), and miss as Δ>10 (i.e., greater than a full-log difference). **(A)** Rift Valley fever virus (RVFV) predictions using manually thresholded images compared to nnU-Net automated segmentation. **(B)** Nipah virus (NiV) predictions using manually thresholded images compared to nnU-Net automated segmentation. **(C)** Severe acute respiratory syndrome coronavirus 2 (SARS-CoV-2) predictions using manually thresholded images compared to nnU-Net automated segmentation. **(D)** Eastern equine encephalitis virus (EEEV) predictions using manually thresholded images compared to nnU-Net automated segmentation.

## Discussion

This study introduced the CPE area assay as a faster, more cost-effective, and less subjective alternative to traditional virus quantification methods. Initial comparisons between the CPE area assay and a traditional six-well plaque assay demonstrated its efficiency and scalability. While the six-well plaque assays required up to 124 plates to process 40 samples, the CPE area assay achieved the same with a single partial 384-well plate. Each 384-well plate can accommodate up to 72 samples, which is equivalent to approximately 216 six-well plates (assuming three plates per sample and excluding controls), highlighting a major improvement in throughput capacity. This increased capacity makes the CPE area assay particularly well-suited for high-throughput screening applications and large-scale studies, offering a substantial improvement in throughput without any considerable loss in data quality.

Optimization of infection conditions was essential in maximizing assay resolution. For each virus tested, multiple time points and a range of virus concentrations were evaluated to determine conditions that produced sufficient CPE development and enabled a robust dynamic range for quantification. Despite optimized conditions, the traditional plaque assay still had a broader quantifiable range, as shown in Fig 3. The traditional plaque assay was able to quantify all samples, regardless of virus concentration, whereas the CPE area assay was unable to quantify anything under approximately 1×10^4^ PFU/mL. Extending the RVFV infection time to 72 h, as illustrated in Fig 2, could have improved the LLOQ. However, this adjustment introduces a tradeoff between broader dynamic range and decreased prediction accuracy at high concentrations. With its higher throughput but smaller dynamic range relative to the plaque assay, the CPE area assay is optimally positioned as a high-throughput screening tool that reliably quantifies samples within range while identifying those outside threshold concentrations.

This study also explored the feasibility and performance of a deep-learning-based approach, particularly CNNs (via SNN) and ViTs, for quantifying CPE in high-throughput viral assays. An SNN was evaluated and, while initial performance on RVFV test dataset was strong at lower concentration range, higher concentrations showed a plateauing effect, either from training or anchor-selection bias. Unlike the CPE area assay, which relied on a 96-well standard curve, the SNN approach used anchor images. This approach offered the theoretical advantage of reducing the number of required control wells to 12, potentially increasing assay throughput. However, this strategy fell short in terms of overall accuracy and needs to be further refined before implemented. Future improvements could involve replacing the current anchor-selection method, which is based on the most generalized mean absolute error, with a method that selects anchors showing the smallest predicted distance or greatest similarity to the experimental samples.

A ViT was fine-tuned on the same datasets as the SNN to evaluate how different deep learning architectures perform in CPE virus quantification. Unlike the SNN, the ViT model predicted continuous values directly from single images without reliance on anchor images. The ViT’s outputs were used to generate four-parameter logistic (4PL) regression models across standard curve values, analogous to the CPE area assay. The ViT predictions closely mirrored biological trends observed in the CPE area assay. This parallel trend suggests that the ViT is either closely aligned with the CPE area signal or learning image features directly associated with it. The model was most precise in the mid-range of the assay, with diminished accuracy at low and high concentration extremes, a limitation also seen in the CPE area assay. Notably, the ViT used 224×224-pixel images due to constraints of the pretrained model architecture, while the CPE area assay used 405×405-pixel images. This reduction in image resolution could partially explain the slight reduction in prediction accuracy for this model compared to the CPE area assay.

One minor limitation in the percent CPE area assay was that it required manual review of the plates and thresholded images. In many cases, thresholds needed to be changed for individual plates for what was considered the “masked” area on the brightfield image and for “positive” on the crystal violet image. To streamline the process of reviewing new plates, we trained an nnU-Net model to segment the images for CPE, cell monolayer, and background using both the crystal violet and brightfield images. This model showed exceptional accuracy in segmenting the image, matching the thresholded images almost perfectly. More importantly, this method was able to quantify samples with at least the same accuracy as the thresholding method, while requiring less effort from the technician running the assay. These methods both showed high accuracy in quantifying virus concentrations of EEEV, NiV, RVFV, and SARS-CoV-2.

## Materials and methods

### Cell and virus maintenance

Grivet (*Chlorocebus aethiops* (Linnaeus, 1758)) Vero (ATCC, #CCL-81) and Vero E6 (BEI Resources, #NR-596) cells were used for virus propagation and maintained in Dulbecco’s Modified Eagle Medium with L-glutamine (DMEM; Gibco, #11995-040), supplemented with 10% heat-inactivated fetal bovine serum (FBS; Sigma Aldrich, #F4135), at 37°C with 5% carbon dioxide (CO_2_). For virus exposure experiments, cells were cultured in DMEM containing 2% heat-inactivated FBS under the same incubation conditions.

All viruses were propagated at the Integrated Research Facility at Fort Detrick in appropriate biosafety containment. The Rift Valley fever virus (RVFV) MP12 isolate was obtained from Dr. Robert Tesh (University of Texas Medical Branch) and propagated in Vero cells and incubated for 72 h under biosafety level 2 (BSL-2) conditions. Eastern equine encephalitis virus (EEEV) FL-93-939 isolate (BEI Resources, #NR-41567), Nipah virus (NiV) Malaysia (United States Army Medical Research Institute of Infectious Diseases), and severe acute respiratory syndrome coronavirus 2 (SARS-CoV-2) WA.01 isolate (U.S. Centers for Disease Control and Prevention) were all propagated in Vero E6 cells under BSL-4 conditions. Nipah virus was incubated for 72 h, whereas EEEV and SARS-CoV-2 were incubated for 48 h. Master stocks for each virus were generated and quantified using six-well plaque assays in Vero E6 cells with a 2.5% microcrystalline cellulose (FMC BioPolymer, RC-591) overlay. After incubation (48 or 72 h, depending on virus), plates were fixed with 10% neutral buffered formalin (NBF; Epredia, #5725) containing 0.2% crystal violet (Ricca Chemical, #3233-16) for plaque visualization.

### Plaque assay (six-well plates)

Vero E6 cells were plated in six-well plates at a density of 1×10^6^ cells per well to reach 90– 100% confluence the following day. Log dilutions were performed on samples in 2-mL deep-well blocks with DMEM containing 10% heat-inactivated FBS. Samples were added, in triplicate of 100 µL per well, to individual wells and incubated at 37°C and 5% CO_2_ for 1 h, with rocking every 15 min. Then, 2 mL of 2.5% microcrystalline cellulose overlay (diluted [1:1] in 2X Modified Eagle Medium [MEM; Gibco, #11935-046] containing 10% heat-inactivated FBS, 2X Antibiotic-Antimycotic [Gibco, #15240-062], and 2X GlutaMAX [Life Technologies, #35050061]) were mixed thoroughly and added to each well. The cells were incubated at 37°C and 5% CO_2_ for 48 h. After incubation, the overlay was removed, and the cells were fixed with 0.2% crystal violet in 10% NBF at room temperature for 30 min. Plates were gently washed with water and the plaques manually counted.

### Cytopathic effect (CPE) area assay

Vero E6 cells were seeded in 384-well plates at a density of 1×10^4^ cells per well in 30 µL of DMEM containing 10% heat-inactivated FBS and incubated overnight at 37°C with 5% CO_2_. The following day, 30 µL of sample or virions were added to each well and incubated for 1 h at 37°C with 5% CO_2_. After incubation, 60 µL of 2.5% microcrystalline cellulose overlay (diluted [1:1] in 2X MEM containing 10% heat-inactivated FBS, 2X Antibiotic-Antimycotic, and 2X GlutaMAX) was added to each well and mixed thoroughly. The cells were incubated at 37°C and 5% CO_2_ for the designated infection period, after which the overlay and media were carefully discarded. Then, cells were fixed using 60 µL of 10% NBF for 1 h at room temperature. Next, NBF was removed and cells were stained with 60 µL of 0.2% crystal violet for 30 min at room temperature. The stain was gently washed off with water, and plates were dried at room temperature overnight prior to imaging on the Operetta CLS high-content imaging system (PerkinElmer/Revvity).

### Cytopathic effect area assay analysis

The Operetta system captured nine images at 10X magnification per well to ensure full coverage of the cell monolayer. Crystal violet images were captured with excitation wavelength of 530– 560 nm and emission wavelength of 655–760 nm. A brightfield image was captured for each well and used to generate a mask that defined the quantification area. These images were exported as TIF files, stitched together to generate a composite image of the entire well, and downsized to 405×405-pixel images to optimize long-term storage. During thresholding of the crystal violet image, only the pixels within the brightfield mask were analyzed, helping to eliminate artifacts and background noise from outside the well. A manual visual check was implemented to review both high- and low-titer images from the standard curve to ensure that the threshold value was appropriately defined. Percent area values from control images were used to construct a standard curve using a four-parameter logistic (4PL) regression function, which was applied to experimental percent area values to calculate final predicted virus concentrations.

### Calculations and statistics

All samples for the CPE area quantification and CPE area reduction assays were run in quadruplicate and passed through a Dixon’s Q-Test at 95% confidence level to remove outliers prior to further analysis [17].

### Siamese neural network training and evaluation

#### Model setup

The Siamese neural network (SNN) model [12, 13] was implemented using the PyTorch library. It comprised two identical convolutional neural networks (CNNs) that were adapted from the standard AlexNet architecture to match the images in the current study. The input number of channels was set to one (for grayscale images) and the final fully connected layer was configured to output a 4,096-dimensional vector. The SNN model used the CNNs to process two 405×405-pixel images simultaneously. Two feature vectors generated by the identical networks are subtracted and the result is passed through a single linear layer to predict the difference in labels (regression value).

#### Training

Ten 384-well plates were exposed to RVFV and fixed at 48 h post-exposure to generate the images for training, validation, and initial test sets. The Operetta imaging system can introduce slight angular deviations across the plate, causing some wells to appear offset; for example, the distance from the left edge of the image to the edge of the well in column 1 differs from that in column 12. To avoid the algorithm associating positional effects with virus concentration, an alternating high-low dilution pattern was adopted. This pattern was applied to five plates and then inverted for the remaining five plates to further reduce positional bias.

The dataset of 3,840 images was stratified and split into a 90:10 ratio, with 90% allocated to training and validation and the remaining 10% reserved for the initial test set. The 90% training/validation set was further split 90:10, resulting in 81% of the total images for training and 9% for validation. The model was trained for 50 epochs using a mean squared error (MSE) loss function to minimize the predicted distance between anchor and experimental images. Anchor images (reference images based on control wells) were selected at random during training and validation.

#### Evaluation

To evaluate the performance of the SNN, a separate test dataset was used in conjunction with a set of pre-selected anchor images. Anchors were selected from a standard curve positioned on the same test plate as the evaluation samples. This standard curve consisted of reference images with known virus concentrations.

Each image in the standard curve was individually treated as a candidate anchor. For a given candidate, the model was used to compute the predicted differences between the candidate anchor and all other images in the standard curve. These predictions were compared to the ground truth differences, and the mean absolute error (MAE) was calculated for each candidate anchor’s prediction set.

This process was repeated for every image in the standard curve, yielding a set of MAE values corresponding to each candidate anchor. The prediction results were then averaged to quantify the overall predictive consistency and effectiveness of each image when used as an anchor. The final anchor set used for test evaluation consisted of those images on the test plate with the lowest average MAE, representing optimal reference points for the model’s comparative assessments.

### Vision transformer

#### Model setup

A ViT model (ViT-B_16) pretrained on ImageNet (1K_V1) was loaded using the torchvision package. To adapt it for regression tasks, the final classification layer was replaced with a linear layer that outputs a single continuous value. All input images were resampled to 224×224 pixels to match the model’s input size requirements.

#### Fine-tuning

The images used for training the SNN were employed to fine-tune the ViT model. The dataset was split into training and validation subsets using an 80/20 split. The split was stratified by the target value (virus concentration) to ensure proportional representation. Hyperparameters were defined for training the model including the number of epochs (10), the image size (224×224 pixels), number of patches (14), learning rate (3×10^-4^), batch size (256), and MSE loss function.

#### Evaluation

The model was evaluated on a separate test dataset that was not used during training. For each input image, the model generated a single output representing the natural logarithm of virus concentration (in PFU/well). Predicted outputs from control wells were used to construct a standard curve using a 4PL regression. This curve was then applied to the model’s predictions for experimental wells to convert them into final estimated virus concentrations.

### nnU-Net segmentation

An nnU-Net segmentation model was trained using grayscale TIF images from both crystal violet and brightfield channels. Images were merged and resampled as described above, with crystal violet assigned to channel 0 and brightfield to channel 1. Default normalization methods provided by nnU-Net were applied to each channel.

Ground truth masks were generated by manually applying intensity thresholds to both the crystal violet and brightfield images. Images were labeled 0 for non-relevant areas, 1 for the CPE area, and 2 for cell monolayer. Thresholds were selected independently for each plate. Areas falling below the brightfield threshold were labeled with 0. Regions above the brightfield threshold were labeled with 1 if below the crystal violet threshold or 2 if above it.

The training set comprised 10,748 images of cells infected with varying doses of EEEV, NiV, RVFV, or SARS-CoV-2. A five-fold cross-validation was performed, and the fold with the highest pseudo-DICE score was selected for downstream analysis. A separate 384-well plate containing EEEV-infected cells at various concentrations and not used during training served as the test set.

## Data availability statement

Code is available at: https://github.com/niaid/irf-area-viral-quant.

Data and images from this study will be made available upon reasonable request.

## Acknowledgments

The authors thank Anya Crane (Integrated Research Facility at Fort Detrick, National Institute of Allergy and Infectious Diseases, National Institutes of Health, Fort Detrick, Frederick, MD, USA) for critically editing the manuscript and Jiro Wada (Integrated Research Facility at Fort Detrick, National Institute of Allergy and Infectious Diseases, National Institutes of Health, Fort Detrick, Frederick, MD, USA) for preparing figures.

## Supporting information

**Fig S1.**
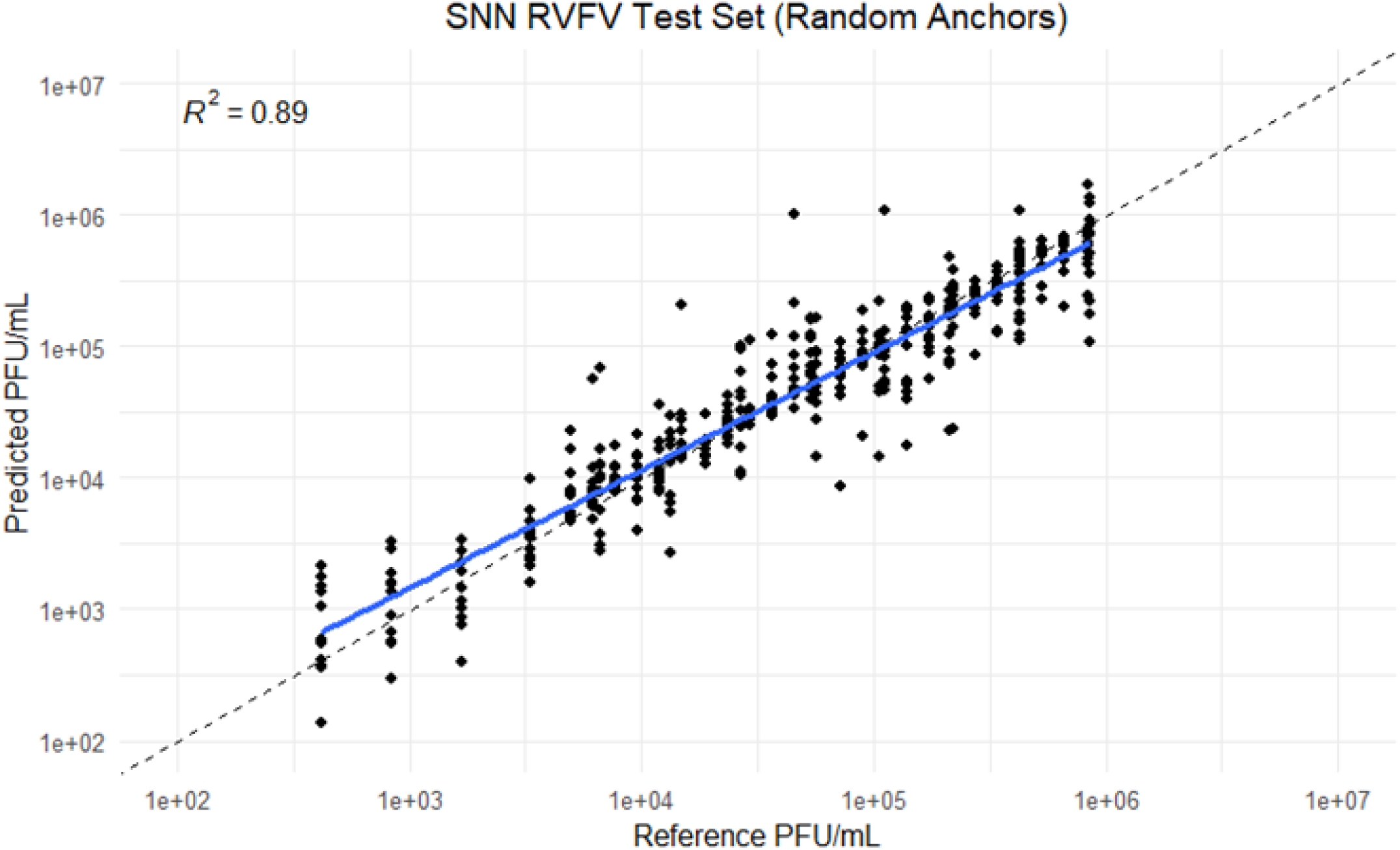
Siamese neural network Rift Valley fever virus test set. Initial test split composed of images from the same plates used in the training and validation sets.

